# WhatsHap: fast and accurate read-based phasing

**DOI:** 10.1101/085050

**Authors:** Marcel Martin, Murray Patterson, Shilpa Garg, Sarah O Fischer, Nadia Pisanti, Gunnar W Klau, Alexander Schöenhuth, Tobias Marschall

## Abstract

Read-based phasing allows to reconstruct the haplotypes of a sample purely from sequencing reads. While phasing is an important step for answering questions about population genetics, compound heterozygosity, and to aid in clinical decision making, there has been a lack of accurate, usable and standards-based software.

WhatsHap is a production-ready tool for highly accurate read-based phasing. It was designed from the beginning to leverage third-generation sequencing technologies, whose long reads can span many variants and are therefore ideal for phasing. WhatsHap works also well with second-generation data, is easy to use and will phase not only SNVs, but also indels and other variants. It is unique in its ability to combine read-based with pedigree-based phasing, allowing to further improve accuracy if multiple related samples are provided.

## 1 Background

Variant calling finds isolated differences between an individual's genome and a known reference. The most common variants are single nucleotide variants (SNVs), insertions and deletions (indels). The set of alleles present at a variant locus is known as *genotype*, such as A/A (homozygous) or G/A (heterozygous) in a diploid individual. Genotype information from a single individual does not allow to draw conclusions about which of the two alleles is paternal and which one is maternal – G/A and A/G are equivalent. Thus, the full sequence of nucleotides in an individual chromosome – the haplotype – cannot be reconstructed. *Phasing* is the process of inferring the correct relationship (*cis or trans*) between alleles at multiple variant loci using additional information besides genotypes, and allows to reconstruct these haplotypes. For example, phasing the two heterozygous variants G/A and C/T may lead to the result that the correct haplotypes are G–T (one copy of the chromosome) and A–C (the other copy of the chromosome).

Phasing allows to obtain crucial insight into various aspects of population genetics, such as population structure, migration and environmental pressure [1, 2]. On an individual level, phased variant data can significantly aid clinical decision making [3]. It is important in investigating compound heterozygosity [4, 2] and allele-specific expression [5].

### 1.1 Approaches

*Molecular phasing* refers to the direct measurement of haplotype sequences. It can be achieved by physically separating the two homologous chromosomes before further processing (such as sequencing), by methods like microscopy-based chromosome Martin et al. Page 2 of 18 isolation, fluorescence-activated sorting, or microfluidics-based sorting [6]. Sequencing haploid cell lines, like CHM1, a hydatidiform mole, can also be considered to fall in this category [7, 8].

*Genetic (pedigree-based) phasing* makes use of family relationships between individuals. For example, if the genotypes at a locus are A/A in the mother, T/T in the father, and A/T in the child, then the A allele of the child is on the maternal haplotype, and the T on the paternal one. Analogous conclusions can be drawn in other cases and in larger pedigrees as long as the variant is homozygous in at least one individual. Refer to the reviews [2, 3] for details and for references to tools that implement these ideas.

*Population-based phasing*, also known as statistical phasing, uses genotype data of large cohorts to infer haplotypes. It is based on the observation that shared ancestry and limited recombination give rise to shared haplotype blocks. This idea is most often formalized in terms of Li-Stephens models [9], with the goal of inferring a maximum-likelihood model given the data. Many different implementations of this idea exist, as reviewed in [1]. These approaches allow either phasing a (large) cohort from genotype data or allow phasing a single additional individual with respect to a haplotype reference panel created previously. The accuracy of such methods crucially depends on the size of the panel and recent progress has been made to scale them up to panels of more than 100 000 haplotypes [10, 11]. However, the decreased performance for rare variants and the inability to phase private and de novo variants remains a limitation.

Finally, *read-based phasing* uses mapped sequencing reads spanning at least two heterozygous variants to infer phase. It is sometimes also called *haplotype assembly*, but should not be confused with haplotype-resolved (*de novo*) genome assembly, which has recently also been explored [12, 13]. Read-based phasing has become increasingly more attractive with the arrival of long-read sequencing technologies because longer reads allow to link more alleles, which yields longer haplotype blocks.

In this paper, we introduce *WhatsHap* as a production-ready and easy-to-use tool for read-based phasing. Its core algorithm for single individuals has been previously described [14]. Beyond that, WhatsHap has the ability to employ pedigree information if provided with multiple, related samples, thus combining read-based and genetic phasing, which is enabled by algorithmic advances previously described [15]. Here, we focus on the WhatsHap tool itself, which includes many additional novel features such as a realignment strategy that significantly enhances phasing performance, support tools to work with haplotype data, as well as phasing of indels and other multi-nucleotide variants.

### 1.2 Tools

Driven by the advances in third-generation sequencing [16, 17, 18] and the corresponding growth in read length, read-based phasing has been gaining considerable attention [19].

Read-based phasing is most commonly formulated as the NP-hard *Minimum Error Correction* problem [20]. To solve it, heuristic [21, 22, 23] and exact approaches [24, 25, 26, 27] have been suggested. In terms of applicable software, the main focus of these approaches has arguably been on proof-of-principle implementations rather han providing production-ready tools. One exception is GATK's [28] ReadBacked-Phasing command, which is one of few tools that directly support standard Variant Call Format (VCF) files for both input and output. However, the underlying algorithm is unpublished and maintenance has been discontinued. HapCUT [22], as another example, is Open Source and even supports phasing of indels, but requires pre-processing of input data and generates non-standard output. Recently, phASER [29] was published with a focus on phasing from RNA sequencing data, but it can also be used for whole genomes. It supports standard VCF output.

**Figure 1.**
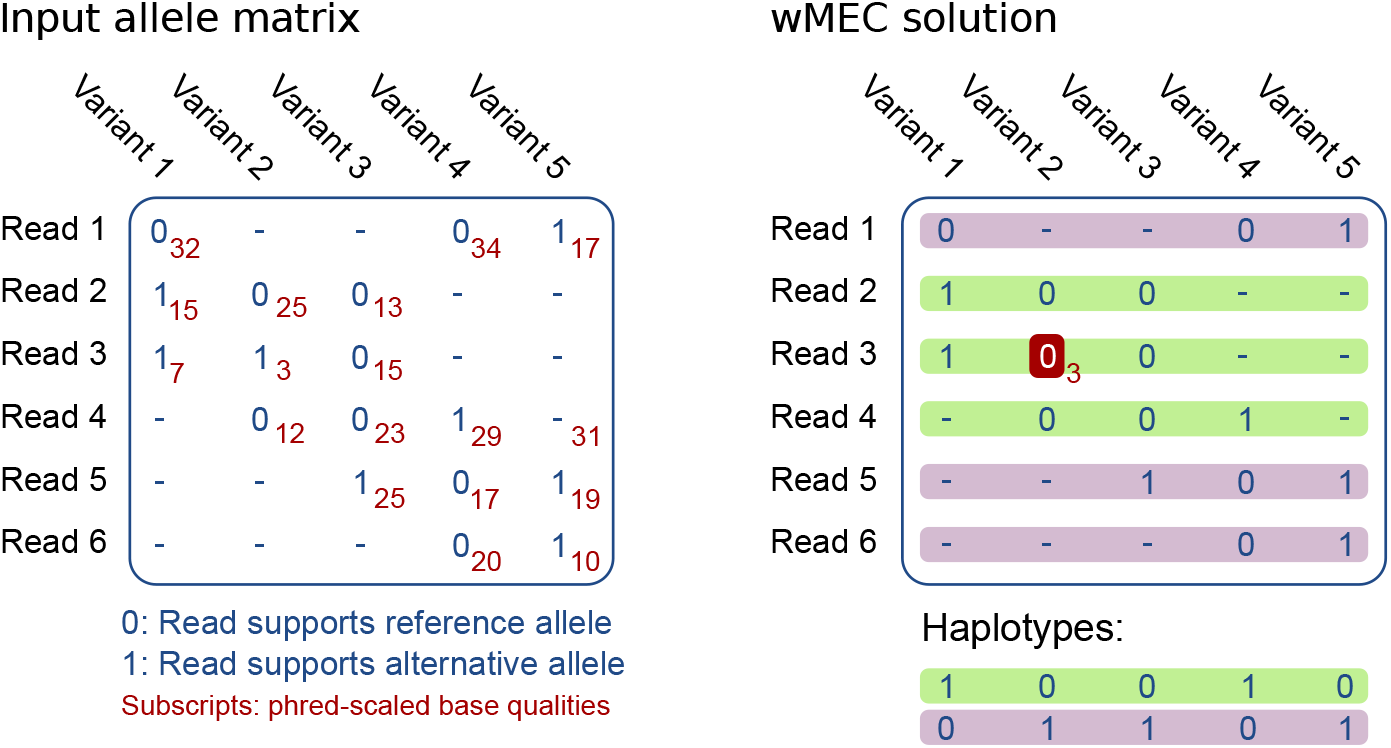
The weighted Minimum Error Correction problem (wMEC) summarized. Left: The allele matrix encodes, for each read, whether it supports the reference (0) or alternative allele (1) at all covered variant positions. Read 1 is a paired-end read. Right: In this example, the solution to the wMEC problem is to flip the single marked entry at Read 3, Variant 2 at a cost of 3. This partitions the set of reads into two conflict-free (internally consistent) sets representing the haplotypes, here color-coded green/purple.

For production use, software is needed that computes high-quality haplotypes, is easy to use, supports standard formats such as VCF and BAM, is well-maintained and does not depend on a particular sequencing technology. WhatsHap meets these requirements while also being based on a solid theoretical framework [14]. It can phase indels, works well with (potentially error-prone) long reads, and, unlike any other tool, can combine read-based and genetic phasing [15]. WhatsHap is available as Open Source software under the MIT license.

## 2 Algorithm and implementation

WhatsHap solves the *weighted minimum error correction problem* (wMEC) for diploid samples. We summarize here the core algorithm from [14].

Conceptually, the algorithm works on an *allele matrix* with one column per heterozygous variant and one row per read, see Figure 1. Allowed entries are 0, 1 and “-”, which denote that the read supports the reference allele, the alternative allele, or that the read does not cover the variant, respectively. For example, rows representing paired-end reads show as two contiguous stretches of 0/1 values, separated by a stretch of “-” at the unsequenced internal segment, as shown in Read 1 in Figure 1. The MEC problem is to correct the smallest number of entries from 0 to 1 or vice versa such that the rows can be partitioned into two haplotypes without conflicts. The wMEC version of the problem takes phred-scaled costs for flipping entries into account.

WhatsHap is an exact, fixed-parameter tractable (FPT) algorithm for the wMEC problem, with read coverage as the only fixed parameter. The algorithm uses dynamic programming to find a partition of the reads into two conflict-free sets that incurs the minimum cost in terms of weights. The algorithms begins at the variant closest to the 5' end of a chromosome and proceeds towards the 3' end along all variant sites. At each site, it computes costs for all 2^k^ possible bipartitions of the k reads covering the respective site. WhatsHap enumerates bipartitions in Gray code order, which allows to compute the cost associated with a particular bipartition in constant time. We refer the reader to [14] for further details on the core algorithm.

The runtime of the algorithm is not affected by the length of the reads. Unlike other phasing tools, WhatsHap is thus well-suited to deal with the continuously increasing read lengths that will arrive with future generations of sequencing technologies.

### 2.1 Detecting alleles in reads

Before the core phasing algorithm can run, the program needs to build the allele matrix from the input data. Variants are expected in standard Variant Call Format (VCF), and aligned sequencing reads are provided as one or multiple BAM files. The input VCF file contains a list of variants and their genotypes. For each read and each heterozygous variant covered by that read, the corresponding entry in the allele matrix needs to be set to 0 if the read supports the reference allele, to 1 if it supports the alternative allele or to “-” if there is no evidence for the variant. Two different strategies are implemented in WhatsHap.

The first is to inspect existing read alignments recorded in the BAM file. Whenever the alignment contains a match to the reference at an SNV position, the allele matrix entry is set to 0, and if it is a mismatch with the appropriate other nucleotide, it is set to 1. For mismatches with one of the other two nucleotides, the entry is set to “-”.

This approach has severe limitations when dealing with raw PacBio or Oxford Nanopore reads, which are affected by error rates of 10% or more. Since this includes indel errors, alignments at variant sites are no longer reliable. As an example, consider the reference sequence TCGTGT with a heterozygous G/A variant at the third nucleotide. One optimal alignment between read TCAGT and the reference could be:

~~~
Read: TC-AGT
Ref.: TC<u>G</u>TGT
~~~

Since the aligner is unaware of the alternative allele, it can place the gap at either pos. 3 or 4 because both options result in the same alignment score (one deletion, one mismatch), resulting in an “Unknown” allele in the shown case. Aligning to the alternative allele instead gives a more correct alignment with a better score (match instead of mismatch):

~~~
Read: TCA-GT
Alt.: TCATGT
~~~

**Figure 2.**
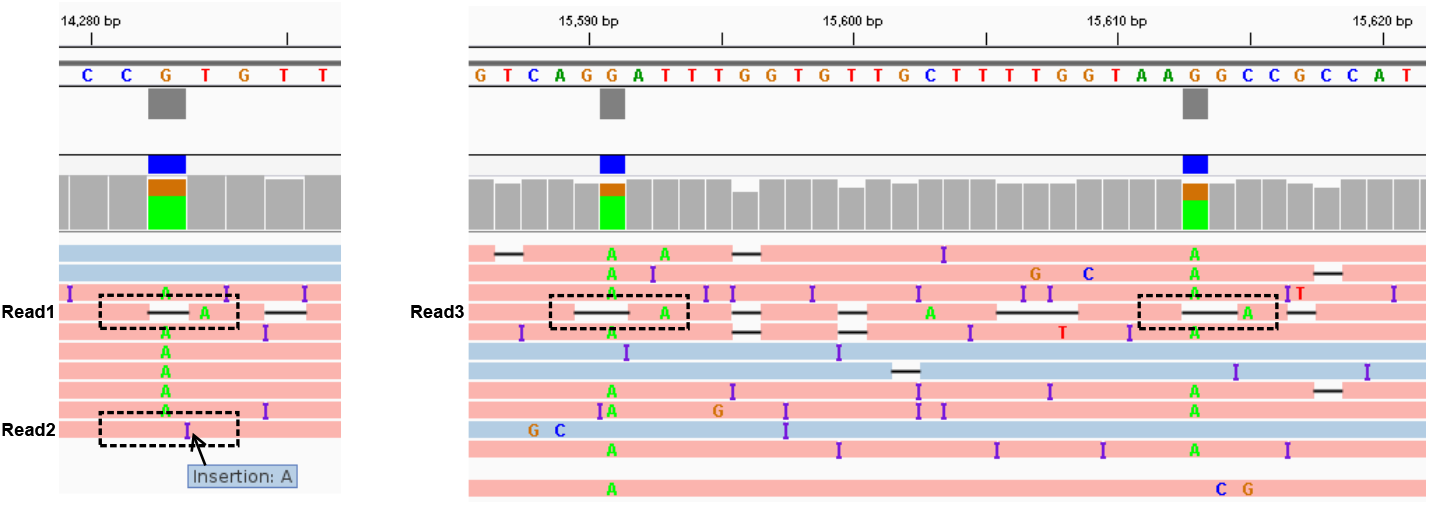
Noisy sequencing data benefits from re-alignment. These screenshots from IGV (Integrative Genomics Viewer) show PacBio reads aligned to three G/A variants. In Read1, the variant is covered by a deletion and would be detected as “unknown”. Re-aligning to the alternative version of the genomic sequence (with an A) would align the read's A on top of the variant, and it would therefore be detected as “alternative”, which is likely the correct explanation. Read2 supports the reference allele although an A insertion is next to the variant. Depending on how alignment scores are chosen, re-alignment allows us to detect either the alternative allele or to determine that both alleles are equally likely. Read3 shows two more examples of alleles that can be detected through re-alignment.

Current aligners are biased towards the reference sequence since they are unaware of the existence of the alternative allele. This leads to misdetected or undetected alleles when the existing read alignment is trusted. Figure 2 illustrates this on real data.

The second allele detection strategy in WhatsHap is therefore based on re-aligning the read in a way similar to what some variant callers do [30]. Using the existing alignment, WhatsHap first extracts the read sequence in a window around the variant, from 10 bp upstream to 10 bp downstream. It then aligns that sequence to the corresponding section in the reference and also to an artificially created sequence that represents what the reference would look like if it incorporated the alternative allele. The allele with a higher alignment score is then assumed to be supported by the read and recorded in the allele matrix; for equal scores, the result is “Unknown”.

This strategy has two advantages: First, it improves the phasing performance for noisy long-read data significantly. Second, we can use the same algorithm to reliably detect and therefore phase indels and multi-nucleotide variants. In fact, any variant specified by providing both a reference and an alternative allele in the input VCF file, with the exception of large structural variants, can be detected by this re-alignment step.

### 2.2 Read selection

Recall that WhatsHap implements a fixed-parameter tractable algorithm with read coverage as fixed parameter. To ensure that read coverage does not exceed a desired threshold, WhatsHap uses a *read selection* heuristic (see [31] for details) as a pre-processing step. The heuristic greedily selects reads that are informative for phasing as long as the coverage threshold is not exceeded. Informative reads are selected based on a per-read score that reflects its suitability for phasing. The score takes into account the number of heterozygous variants it covers and the number of “holes”, that is, the number of “-” entries that are physically covered by the read and hence lie between the first and the last non-dash entry. Such holes are penalized because they do contribute to the physical coverage at a variant locus while not providing information on its phase. To break ties, mapping and base-calling quality are considered. In addition to greedily selecting high-scoring reads, low-scoring reads are added that allow to “bridge” islands of otherwise unconnected variants. Examples of such reads are mate pairs, which span distant variant sites but are otherwise not chosen because their ends cover too few variants and they exhibit many holes.

It was previously demonstrated that this read selection strategy leads to superior results in comparison to random downsampling [31]. It is particularly powerful when feeding WhatsHap with sequencing libraries of mixed origin, such as combining PacBio and Illumina reads. By default, read selection reduces coverage to 15×. Earlier experiments have shown that exceeding 15× coverage mostly leads to adding redundant information [32].

### 2.3 Software engineering

The command-line front-end of WhatsHap is written in Python 3 due to its simplicity and maintainability. The core dynamic programming algorithm that solves the wMEC problem is implemented in C++. The interface between Python and C++ is written in Cython [33], which is a Python dialect that compiles directly to C/C++. Other performance-critical parts have been implemented directly in Cython, such as the re-alignment and read selection algorithms.

External libraries used include PyVCF [34], pysam [35] and pyfaidx [36].

### 2.4 Usage and file formats

WhatsHap is easily installed from the *Python Package Index* (PyPI) or Conda's bioconda channel [37] with a single `pip3 install or conda install` command.

WhatsHap is a command-line tool that expects– in the common case– a single Variant Call Format (VCF) file and one or more BAM files as input. The command `whatshap phase–o phased.vcf input.vcf input.bam` is typically sufficient. Instead of BAM, phasing information can be provided to WhatsHap also as phased VCF files, such as those generated by 10X Genomics' phasing pipeline, and already phased blocks are then treated as “reads”. WhatsHap phases all samples it finds in a multi-sample VCF, using BAM read groups and VCF sample names to associate each read to its originating sample.

Input reads– even those from a single sample– can be distributed over several BAM (or phased VCF) files, and may use different sequencing technologies. For example, PacBio, Illumina mate-pair reads, and 10X Genomics phased blocks can be used simultaneously for phasing.

WhatsHap outputs an augmented version of the input VCF file, where phasing information for those variants that could be phased has been added. By default, phase information is written to the “GT” (genotype) and “PS” (phase set) tags as standardized in the VCF specification. The “HP” tag– compatible with GATK ReadBackedPhasing output– is also supported.

### 2.5 Combining read-based and genetic phasing

The core algorithm for solving the wMEC problem [14] and most of the features we have outlined thus far involve the phasing of a single individual. Recently, Garg et al. [15] have extended the wMEC formalization to exploit pedigree information between a set of related individuals, referred to as *PedMEC*.

Given multiple samples, the corresponding reads, and a pedigree describing the relationship between samples, a solution to the PedMEC problem minimizes error corrections (like wMEC) in such a way that the Mendelian rules of inheritance imposed by the pedigree are respected and that a minimum recombination cost is incurred.

Pedigree-aware phasing thus makes use of the fact that the haplotypes of a child are a recombination of the parental haplotypes. Combining read-based and genetic phasing is principally more powerful than either method alone: Read-based phasing is limited by read length while genetic phasing, on the other hand, cannot phase variants that are heterozygous in all samples, and it cannot find recombination breakpoints in trios.

Recall that WhatsHap can phase multiple samples that are not related if provided with a multi-sample VCF file and appropriate reads in one or more BAM or phased VCF files. WhatsHap switches to the combined read-based/pedigree phasing algorithm if it is additionally supplied with a PED file [38] that describes the relationship between the samples. Recombination events are by default assumed to be equally likely at any position. For more realistic modeling of, for example, recombination hotspots, a vector of recombination rates along the genome (genetic map) can optionally be given.

We have demonstrated earlier [15] that adding pedigree information boosts accuracy immensely. In some cases, the accuracy attained with 15× coverage on a single individual can be achieved with just 2× coverage when a mother-father-child trio is provided.

When in pedigree mode, even variants that are not connected by any read in any sample can often be connected correctly based on pedigree information, allowing to obtain chromosome-length haplotypes when phasing families. In case no read data is available, and hence no BAM files are provided on the command-line, WhatsHap still solves the PedMEC problem and, by doing this, computes a purely pedigree-based phasing.

### 2.6 Supporting tools

Beyond phasing, WhatsHap comes with a collection of other functionalities to handle phasing data. These functionalities are available as sub-commands, allowing to conveniently combine them into larger analysis pipelines. Available sub-commands include “unphase” to remove all phasing information present in a VCF file; “stats” to compute statistics on a phased VCF such as the number of (un)phased variants, number of phased blocks, and minimum/maximum/average block size; and “compare” to compare two or more phased VCFs with respect to pair-wise switch/ flip errors, but also with respect to a simultaneous comparison of multiple phasings. The sub-command “haplotag” uses a phased VCF to add tags to each read in a BAM file that indicate which haplotype it belongs to. This allows further processing, such as splitting the reads into two BAM files according to haplotype, or to color-code reads by haplotype in a genome browser (as for example seen in Figure 2).

## 3 Results

### 3.1 Experimental setup

We evaluate WhatsHap and other phasing tools on both real and simulated datasets. The setup we explain in the following is similar to the one described earlier [15].

*Real data*. As evaluation sample, we chose the child (NA24385) of the Ashkenazim trio sequenced by the Genome in a Bottle Consortium (GIAB) [39]. We downloaded a consensus genotype call set (NIST CallsIn2Technologies 05182015) for chromosome 1, containing 48 023 heterozygous SNVs and 4 265 heterozygous non-SNVs. We phased all these variants with the population-based phasing tool SHAPEIT (v2-r837) [40] with default parameters, using the reference panel provided by Phase 3 of the 1000 Genomes Project. This statistically phased dataset serves as a reference to which we compare results from the read-based phasing tools. Furthermore, we use the Pacific Biosciences (PacBio) reads from GIAB aligned to chromosome 1. They have an average coverage of 60.2× and average mapped read-length of 8 687 bp. To analyze the effect of sequencing depth on phasing performance, we created additional datasets by randomly downsampling the aligned reads to average coverages of 2 ×, 3 ×, 4×, 5×, 10× and 15 ×.

*Simulated data*. While population-based phasing is very accurate for large cohorts, it has limitations with respect to low-frequency variants. Thus, differences between statistical and read-based phasing may be caused by errors in either method. For a more accurate picture of the performance of read-based phasing alone, we complement the evaluation with a simulation in which the true haplotypes are known.

To this end, we generated two haplotypes of a “virtual child” by recombining the haplotypes of the parents NA24143 and NA24149 *in silico* as described previously [15]. The resulting virtual child has 45 625 heterozygous variants with known phasing, containing 42 062 SNVs and 3 563 non-SNVs. Next, we incorporated these true variants into the GRCh37 reference genome to obtain two FASTA files representing the true haplotypes. From those, we simulated reads that mimic the read length profile and coverage of the real data using pbsim [41]. The resulting reads were then aligned to GRCh37 using BWA-MEM (0.7.12-r1039) [42]. As for the real dataset, we further downsampled the aligned reads to average coverages of 2 ×, 3×, 4 ×, 5 ×, 10×, and 15 ×.

### 3.2 Tools

We compared WhatsHap to other phasing approaches that had an implementation available. In addition to WhatsHap, we included the ReadBackedPhasing component of the Genome Analysis Toolkit (GATK) [28]), hapCUT [22] and phASER [29]. For the criteria of how we selected these and notes on alternative programs, [43, 44, 23, 45], see Section 1 in the Supplement,

We used ReadBackedPhasing as distributed with GATK v3.5-0-g36282e4, Git revision 844af08c of hapCUT (reporting as version 0.7), revision 8608e1d of phASER (version 0.8) with a fix for CIGAR hard-clipping available in pull request #14. All three programs required either modifications to the source code or special parameters to work with PacBio data (see Supplement). We used revision 09326a5 (version 0.12) of WhatsHap.

ReadBackedPhasing did not phase any variants in PacBio data with default settings, and there is no official recommendation for the correct settings. We therefore tried 144 parameter combinations on a subset of the data and determined that using options–`phaseQualityThresh 1 and–min base quality score 1` gives the best results (see Supplement).

For hapCUT, a separate program needs to be run that pre-processes BAM/VCF input data. Runtimes that we report later include this step. We added a third step, in which the custom output format is converted to VCF. We did not use hapCUT's–`longreads` option since the help states that it should be set if the “program uses too much memory”, which was not the case.

PhASER was primarily designed for phasing RNA-seq data. Thus, its performance and runtime in our evaluation may not be representative of what it would achieve in an RNA-seq setting. PhASER does not require pre–or postprocessing steps.

We ran all programs both with and without requesting phasing of non-SNVs (mostly indels), except ReadBackedPhasing, which does not support indel phasing. WhatsHap was additionally run in a non-realignment mode, in which it also does not support non-SNV phasing.

### 3.3 Measuring performance

We describe performance metrics that we use to evaluate *phasing accuracy* and *phasing completeness* of the different tools.

For phasing completeness, we need to consider that read-based phasing requires reads covering two heterozygous variants. The result is therefore a partition of the list of variants into *haplotype blocks*, within which all of the variants are phased relative to each other. The theoretically best possible case is that these blocks correspond to the connected components of a connectivity graph, which is a graph with heterozygous variants as the nodes and where an edge is drawn between two variants if a read exists that covers both. In practice, different tools may choose to not use some reads or may have different notions of connectivity and will therefore output haplotype blocks that are more fragmented. A haplotype block with only a single variant is a *singleton*.

In a VCF file, haplotype blocks are called *phase* sets and have a unique identifier, which is attached to a genotype using the PS tag.

In a block of size *k*, we say that there are *k*-1 *phase connections* between adjacent variants, assuming the variants are sorted by position. In our evaluation, we count phase connections because they take the block structure into account; the number of phase connections is equal to the number of phased variants minus the number of blocks.

In our results, we will give the *phase connection ratio*, which is the total number of phase connections divided by the possible number of phase connections (*n*-1 for a chromosome with *n* heterozygous variants). For a read-based phaser, the phase connection ratio can in principle not exceed the theoretical optimum 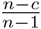, where c is the number of connected components in the connectivity graph of the input data.

**Figure 3.**
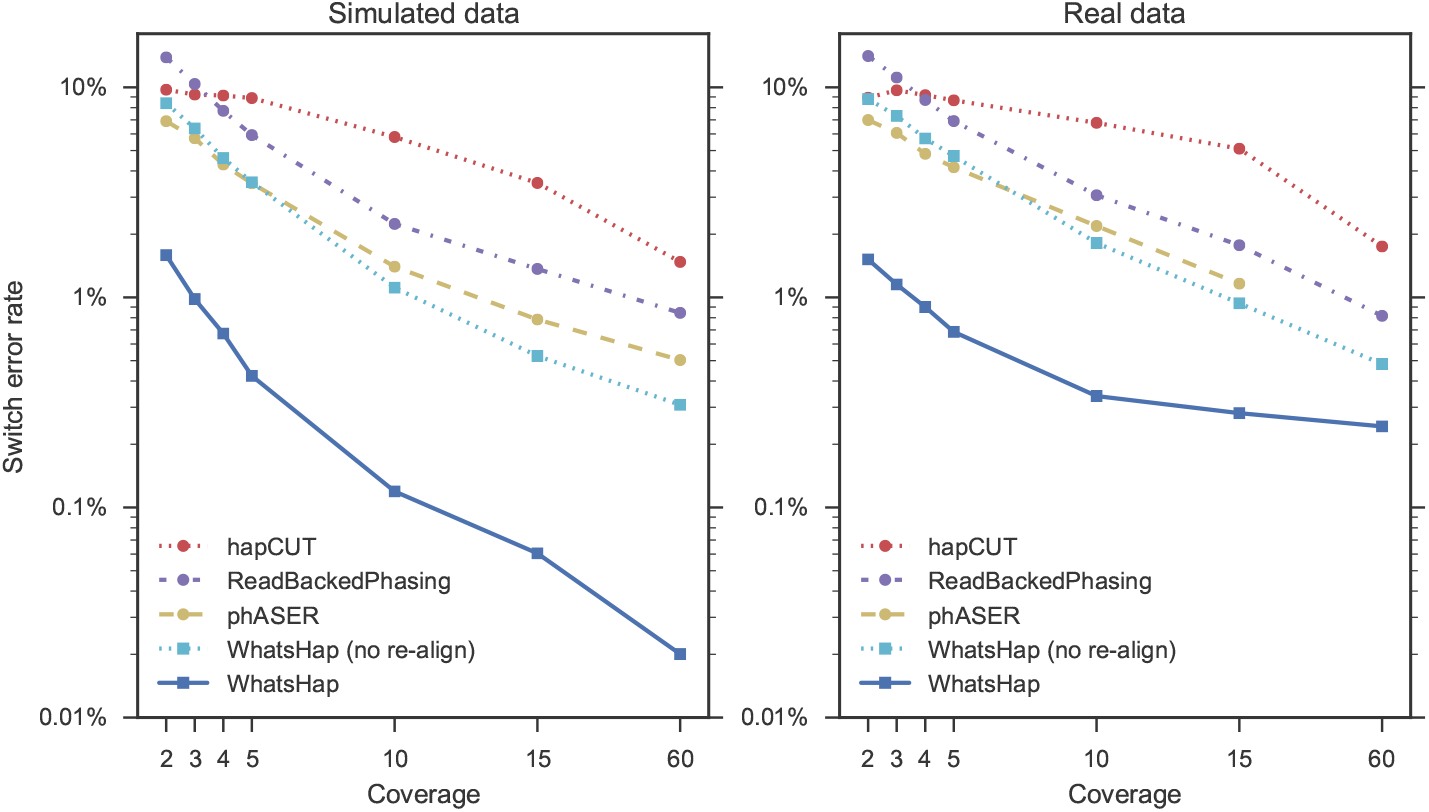
Phasing switch errors on simulated and real PacBio data, SNVs only. All tools give fewer errors with increasing coverage. phASER at full coverage on real data is not shown since the run did not complete within 24 hours. Note logarithmic scaling of the y axis, which emphasizes the difficulty of lowering the error rate the closer it already is to zero.

*Phase* can be seen as a relationship between variants along a phase connection: It can be either *cis*, meaning that the reference alleles of the connected variants are on the same haplotype, or it can be trans, meaning that the reference alleles are on different haplotypes.

Errors within phased blocks are counted in terms of *switch errors*, which occur when the meaning of cis/trans is reversed from one point on forward within a block. Switch errors are calculated by traversing the haplotype blocks from left to right and computing the number of times a jump is needed from one haplotype to the other in order to reconstruct the true haplotype. A flip error, defined as an isolated variant assigned to the wrong haplotype, is counted as two switch errors in this model. Refer to Figure 5a for an example. For the *phasing error rate*, we sum the switch errors over all blocks and divide by the total number of phase connections. Switch errors are detected by comparison to statistical phasing for real data and to the known truth for simulated data.

### 3.4 Results on SNVs only

As one would expect, the five evaluated algorithms consistently perform better when coverage increases. For both simulated and real data, switch error rate decreases (Figure 3) and the phase connection ratio increases (Figure 4).

WhatsHap with the re-alignment algorithm is clearly the tool with the lowest switch error rate at all coverages. In real data and at full coverage, it achieves a switch error rate of 0.24%. The version of WhatsHap without realignment reaches 0.48% at full coverage. While the phASER run for full coverage did not finish within 24 hours, the trend suggests that it could possibly also achieve a rate of around 0.5% in this dataset.

For simulated data, the shapes of the curves and the resulting ranking of tools are mostly the same as for real data, suggesting that the simulation pipeline mimics realistic conditions. WhatsHap achieves an extremely low error rate of 0.02% at full coverage, corresponding to just eight switch errors. WhatsHap without re-alignment achieves 0.3% switch error rate at full coverage in simulated data.

**Figure 4.**
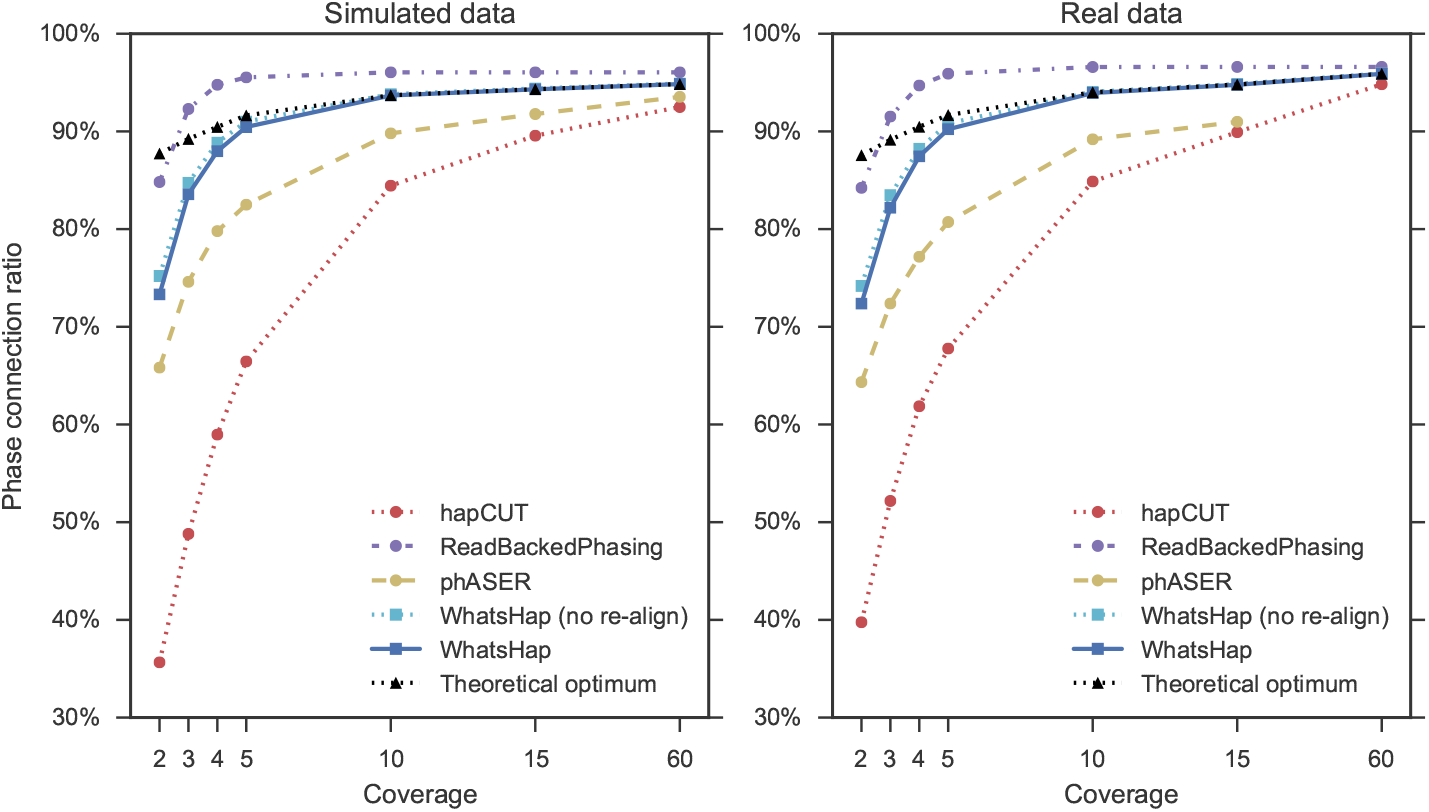
Completeness of phasing on simulated and real PacBio data, SNVs only. All tools make more phase connections as coverage increases. phASER at full coverage on real data is not shown since the run did not complete within 24 hours. The results for ReadBackedPhasing are higher than the theoretical optimum given the reads, see main text.

As a notable exception where the switch error curves for real and simulated data do not agree well, WhatsHap's error rate decreases almost linearly (in this log-scale plot) for simulated data, while reaching a plateau for real data at a level slightly above 0.2%. Recall that phasing errors in real data are computed by comparison to statistical phasing, which is not perfect and makes errors at rare variants. Therefore, some “baseline” errors remaining for all tools in real data may actually be explained by errors made by the statistical phaser.

Figure 4 shows the phase connection ratio the tools were able to achieve. All tools, as expected, connect more variants when coverage increases. At first sight, ReadBackedPhasing seems to perform best as it connects more variants than the other tools. However, for coverages 3× and higher, its phase connection ratio is actually higher than the theoretical optimum. In other words, ReadBackedPhasing outputs *fewer* blocks than there are connected components, which in turn means that it connects variants that are in fact not connected by reads. By inspecting the haplotype blocks it outputs together with the reads and variants in IGV, we could confirm this unexpected behavior, see Supplement for an example.

The phase connection ratio achieved by WhatsHap is close to the optimum, which is to be expected as the haplotype blocks it outputs are constructed from the connected components of the selected reads.

We would like to emphasize the profound impact of our realignment strategy on the results. While the completeness of phasing is virtually the same when WhatsHap is run with and without realignment (Figure 4), the switch error rates improve drastically (Figure 3). For instance, for 10-fold coverage, they decrease from 1.1% to 0.1% on simulated data and from 1.8% to 0.3% on real data. Figure 3 clearly shows that the effect of this novel strategy dwarfs the differences that are due to different phasing algorithms.

**Figure 5.**
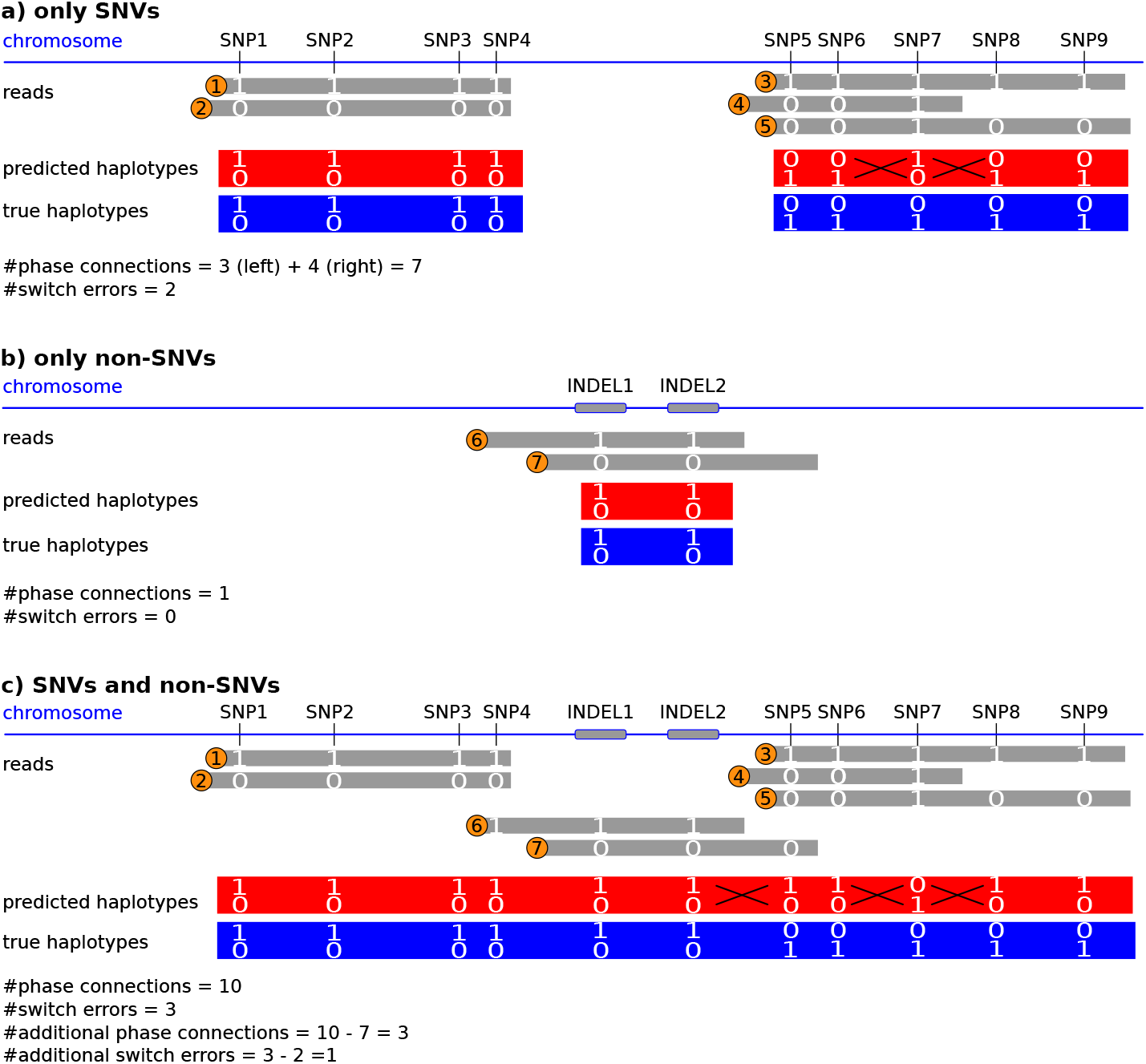
Integrative phasing of SNVs and non-SNVs. Heterozygous variants are shown along a chromosome, together with reads covering them (grey bars) and resulting phased blocks (red bars). Numbers in orange circles indicate read identifiers to emphasize that the same read set is used across panels a), b) and c). For each of the three panels, the number of phase connections is given, which for a single block is its number of variants minus 1. Switch errors are marked by black crosses. Examples are shown when a) only phasing SNVs, b) when only phasing indels and c) when using both simultaneously (bottom).

### 3.5 Results on non-SNVs

WhatsHap, hapCUT and phASER are able to phase insertions and deletions. WhatsHap additionally phases “complex” variants such as CGA → CTCC that are neither SNV, insertion nor deletion. WhatsHap can, in principle, phase all variants whose reference and alternative alleles are given explicitly and are shorter than the read length.

To assess how the three tools handle non-SNVs, we compare the results for phasing only SNVs to the results for jointly phasing both SNVs and non-SNVs. Figure 5 illustrates that we not only expect to see additionally phased non-SNVs, but also an increase in the number of phased SNVs as previously unconnected blocks can become connected due to reads that cover SNVs as well as non-SNVs. Note that phasing only non-SNVs is not useful for assessing performance as this will lead to an extremely fragmented phasing (even long reads rarely cover more than one non-SNV).

**Figure 6.**
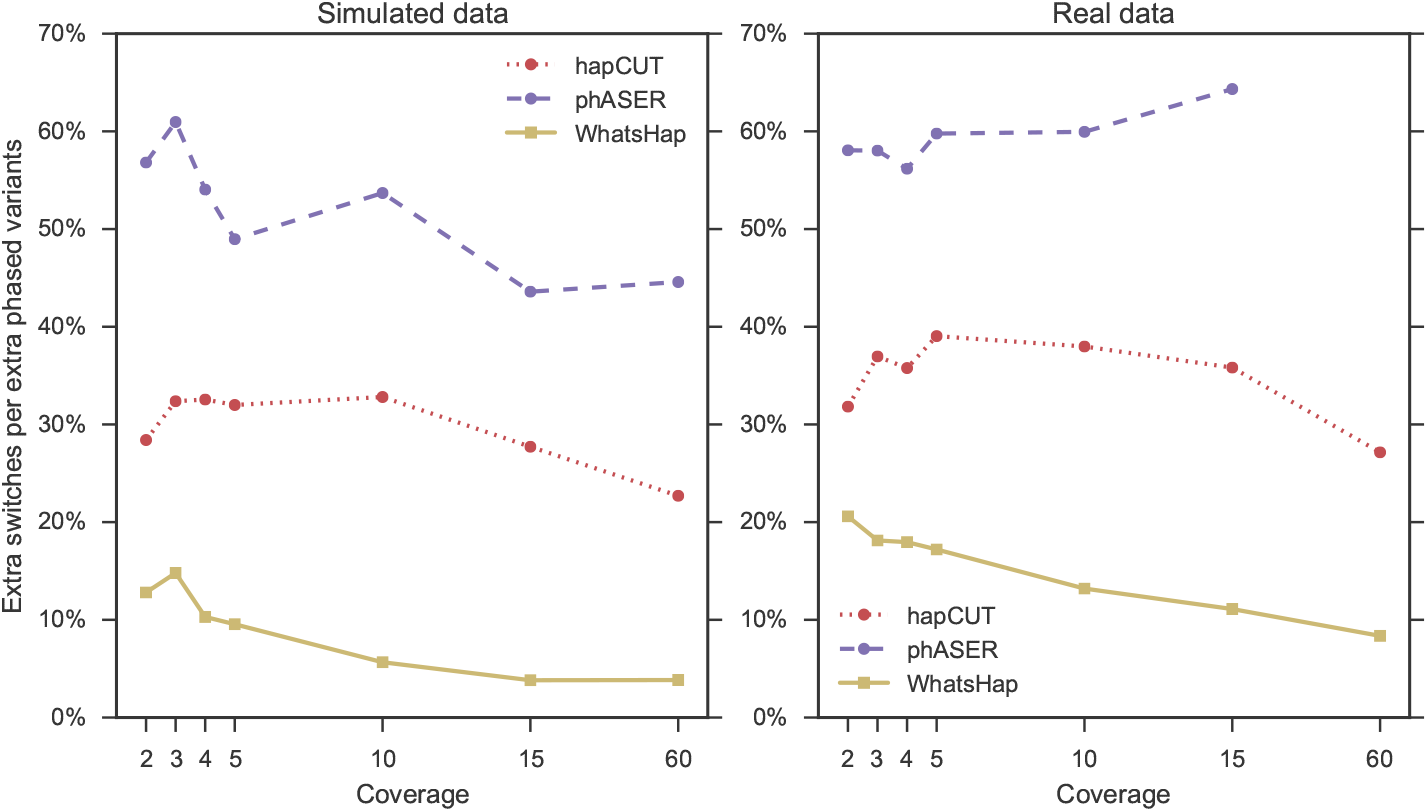
Additional switch errors when phasing non-SNVs. Enabling phasing of non-SNVs, the number of phased variants increases (Δp), but so does the number of switch errors (Δe). The y axis shows 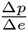.

When phasing of non-SNVs is enabled, the number of phased variants increases (Δp), but so does the number of switch errors (Δe). We define the switch error rate over additionally phased variants as the number of additional switch errors divided by the number of additionally phased variants, 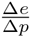. The quantity can be interpreted similar to a phasing error rate, but it takes into account not only switch errors among the phased non-SNVs, but even additional errors among SNVs. For example, a rate of 10% means that ten additionally phased variants come at a price of one additional switch error.

As we can see in Figure 6, the switch error rate over additionally phased variants is in general much higher than the switch error rate in the SNV-only case. Regardless of coverage, HapCUT and phASER are at a rate of over 20%. That is, every fifth additionally phased variant introduces an additional switch error.

WhatsHap is clearly the best of the tools, achieving a rate of 3.8% in simulated data and 8.3% in real data at full coverage. As mentioned before, part of the differences between the read-based phasings and the statistical phasing on the real data might be explained by errors in the statistical phasing. Since indels are difficult to genotype accurately, it seems likely that statistical phasers will exhibit higher error rates for indels than for SNVs. However, even on simulated data the switch error rates in the SNV-only case are at least an order of magnitude lower, which indicates that phasing of non-SNVs is more difficult than phasing SNVs.

Finally, Figure 7 shows how many non-SNVs could actually be phased by the programs when instructed to do so. WhatsHap is again the best choice at any coverage. At full coverage, WhatsHap is able to phase 97% of all non-SNVs in both simulated and real data, while none of the other tools reach more than 90% phased non-SNVs.

**Figure 7.**
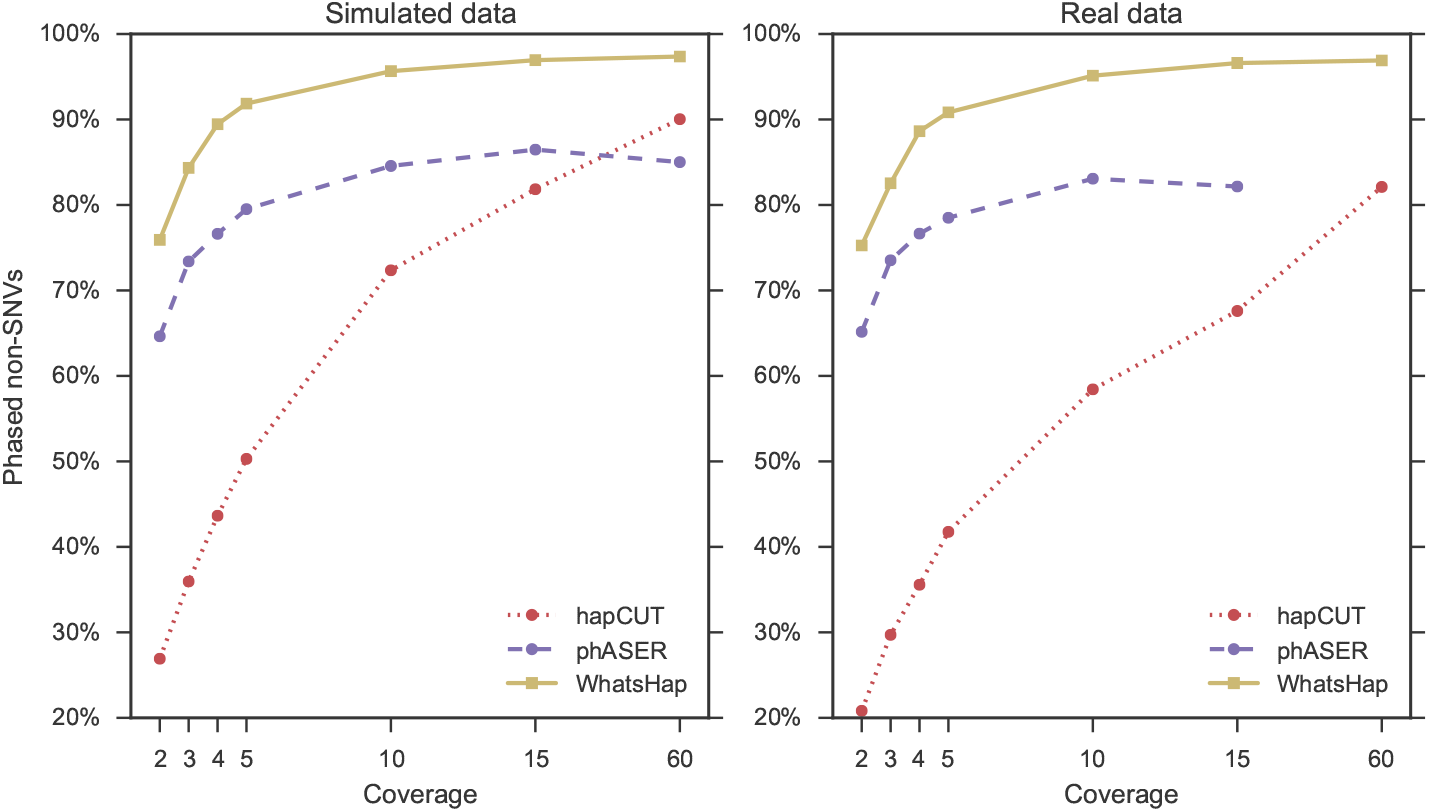
Fraction of non-SNVs that can be phased. The y axis shows how many non-SNVs were phased by each relative to the total number of heterozygous non-SNVs in the dataset.

**Table 1.**
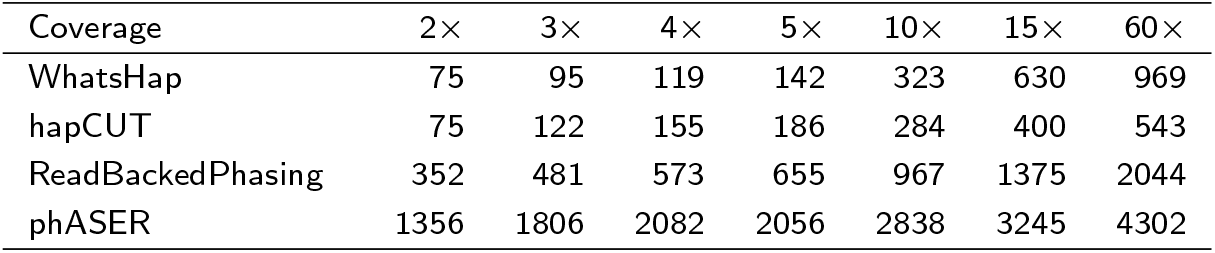
Running time of phasing tools on human chromosome 1, measured on the simulated child dataset. Values given are in CPU seconds; actual (wall-clock) times are higher due to I/O. Note that phASER runtimes includes computations that are applicable only in an RNA-seq setting. The times displayed for WhatsHap are measured with re-alignment mode enabled.

| Coverage | 2× | 3× | 4× | 5× | 10× | 15× | 60× |
| --- | --- | --- | --- | --- | --- | --- | --- |
| WhatsHap | 75 | 95 | 119 | 142 | 323 | 630 | 969 |
| hapCUT | 75 | 122 | 155 | 186 | 284 | 400 | 543 |
| ReadBackedPhasing | 352 | 481 | 573 | 655 | 967 | 1375 | 2044 |
| phASER | 1356 | 1806 | 2082 | 2056 | 2838 | 3245 | 4302 |

### 3.6 Runtime

We measured running time of all tools on the simulated dataset. To be able to include all tools, only SNVs were phased. For those tools supporting non-SNV phasing, the rankings are identical with only slightly higher runtimes (not shown). Results can be seen in Table 1.

WhatsHap's runtime is clearly competitive with those of the other tools. At all coverages, either WhatsHap or hapCUT is the fastest tool. HapCUT has an advantage at high coverages, while WhatsHap is faster at low coverages.

## 4 Discussion and conclusions

The benefits of long reads for the purpose of read-based phasing are obvious. However, most existing tools have been designed without taking the algorithmic challenges of long reads, namely their length and their elevated error rates, into account. WhatsHap is a production-ready tool for read-based phasing that addresses these challenges. WhatsHap's haplotypes are closest to the theoretical optimum in terms of completeness and most accurate in terms of phasing errors incurred, together showing superior performance across all relevant coverages in comparison to other tools. WhatsHap also proves to be superior in terms of phasing non-SNVs, clearly a challenge for all phasing tools.

WhatsHap's core algorithm is based on a sound theoretical framework [14, 15] that delivers optimal solutions to the wMEC or PedMEC problem for bounded coverage. The combination of our exact core algorithm with a heuristic read selection procedure results in an implementation that is fast and accurate in practice. Indeed we observed that runtimes are dominated by I/O operations, not by the phasing algorithm itself.

The program's ability to integrate read-based and pedigree phasing is unique and is the perfect match for studies in which family trios are sequenced. As shown earlier [15], the phasing results obtained in such a setting outperform non-integrated approaches, while reducing the needed overall coverage to as low as 2 ×.

Most importantly, we also consider WhatsHap to be the tool that is easiest to use. Options are set to sensible defaults and typically do not need to be changed by the user. Input formats are recognized automatically, and output is standards-compliant VCF, ready to be consumed by other tools. The ability of using phased VCFs as input allows interesting use cases such as “merging” phasing results. In fact, reads from any sequencing technology and previously-phased data are supported, allowing to combine data from Illumina (paired-end and mate-pairs), PacBio, Oxford Nanopore, 10X Genomics and Hi-C at will.

WhatsHap additionally comes with a set of extra tools that help in analyzing phasing results, such as obtaining statistics from phased VCFs or comparing them.

### 4.1 Future work

WhatsHap is being improved continually, especially in terms of performance, speed and the ability to work with a wide range of types of input data. Beyond that, we aim to integrate population-based phasing capabilities. In the same way that genetic phasing helps to bridge regions insufficiently covered by reads in related samples, population-based phasing can help to achieve this even in single samples given a reference panel. Over time, we predict that phasing tools will necessarily have to focus more on these integrative approaches that make best use of all available data.

We also plan to enrich WhatsHap with other algorithmic backends, in addition to the core phasing algorithm. In particular, HapCol [46] is an alternative exact FPT algorithm for the wMEC problem, where the only fixed parameter is the maximum number of corrections k needed to the set of entries at each variant position, such that the reads can be partitioned into two haplotypes without conflicts. Combining this with ideas from [47], we plan to deliver a method that can do phasing on trios (or larger pedigrees). This will be a promising alternative to PedMEC in situations where this k is much smaller than the coverage. Beyond FPT approaches, work on integrating an integer-linear programming solver for wMEC is underway. In this way, we aim to always provide the best possible solver for a problem instance at hand and, at the same time, provide a test bed for benchmarking new algorithms.

### 4.2 Software availability

WhatsHap is available as Open Source software under the MIT license. The home page at https://whatshap.readthedocs.io/ contains documentation and installation instructions. Supported operating systems include Linux and OS X. Installation requires Python 3.3 or later and a working C/C++ compiler.

All files and instructions necessary to reproduce the results of this study are available at https://bitbucket.org/whatshap/phasing-comparison-experiments/.

## Author's contributions

MP, TM, NP, GK and AS co-invented the WhatsHap algorithm. MM implemented a major part of WhatsHap's Python frontend, implemented the re-alignment algorithm and ran the evaluation. MP and TM implemented the core WhatsHap algorithm. SG and TM contributed the pedigree phasing mode. SOF and TM contributed the read selection algorithm. MM, MP, TM and SG wrote the manuscript, with help from all authors. All authors read and approved the final manuscript.

## Acknowledgements

MM is supported by a grant from the Swedish Knut and Alice Wallenberg Foundation to the Wallenberg Advanced Bioinformatics Infrastructure. AS acknowledges funding from NWO through Vidi grant 639.072.309.

